# Closing the genome of *Teredinibacter turnerae* T7902 by long-read nanopore sequencing

**DOI:** 10.1101/2024.07.30.605897

**Authors:** Mark T. Gasser, Annie Liu, Ron Flatau, Marvin Altamia, Claire Marie Filone, Dan Distel

**Author notes:** Corresponding author DLD.

## Abstract

We present the complete closed circular genome sequence derived from Oxford Nanopore sequencing of the shipworm endosymbiont *Teredinibacter turnerae* T7902 (DSM 15152, ATCC 39867), originally isolated from the shipworm *Lyrodus pedicellatus* (1). This sequence will aid in the comparative genomics of shipworm endosymbionts and the understanding of host-symbiont evolution.

## Announcement

*Teredinibacter* species are cellulolytic gammaproteobacteria (Cellvibrionaceae) that occur as intracellular endosymbionts of wood-boring bivalves (Teredinidae) (1–4), commonly known as shipworms. Strain T7902 was isolated from the gills of a single specimen of the shipworm *Lyrodus pedicellatus* collected in Long Beach, CA, in 1979 and was the second strain of *T. turnerae* brought into pure culture (1). It is the original representative of *T. turnerae* Clade II, one of two distinct clades previously identified among *T. turnerae* strains (5, 6). Gills were dissected, washed in sterile seawater, and homogenized in shipworm basal medium (SBM) (7), and the homogenate was streaked onto a culture plate containing 0.9% agar and SBM at pH 8.0 supplemented with 0.2% w/v powdered cellulose (Sigmacell Type 100; Sigma-Aldrich) and 5 m*M* ammonium chloride. Individual colonies were picked and restreaked on fresh plates until a pure clonal isolate was obtained. The original genome sequence of *T*.*turnerae* T7902 was published to GenBank (GCA_000379165.1) but has not been described previously in peer-reviewed literature. This sequence was completed on 2012-05-22 at the DOE Joint Genome Institute under award 10.46936/10.25585/60001419 using 454 GS FLX Titanium and Illumina HiSeq 2000 sequencing platforms. It was assembled using Velvet v. 1.0.13 (8) and ALLPATHS v. R40295 (9), resulting in an improved high-quality draft assembly comprised of 72 scaffolds with 76 contigs. At the time of this work, the genomes of nine strains of *T. turnerae* were publicly available at the National Center for Biotechnology Information (NCBI), and only one strain genome, T7901 (Clade I, Genbank: GCA_000023025.1), was complete and closed. Here, we present the re-sequencing and completed genome of strain T7902 from nanopore-only sequencing (Table 1).

**Table 1.** Assemblies of Teredinibacter turnerae T7902

| GenBank Assembly | Scaffolds (contigs) | Size (bp) | GC% | CDS |
| --- | --- | --- | --- | --- |
| GCA_000379165.1 | 72 (76) | 5,387,817 | 50.8 | 4268 |
| GCA_037935975.1 (this work) | 1 (1) | 5,348,823 | 50.9 | 4212 |

*Teredinibacter turnerae* strain T7902 colonies were maintained at 30°C on shipworm basal medium (SBM) (7) plates supplemented with 0.025% NH_4_Cl and 0.2% cellulose (Sigmacell Type 101; Sigma-Aldrich). A colony was selected and used to inoculate a 6 mL liquid culture of SBM supplemented with 0.025% NH_4_Cl and 0.2% carboxymethyl cellulose (CMC) medium and grown at 30°C, 100 rpm for 4 days. Bacterial cells were harvested by centrifugation (10 minutes, 4°C, 4,000 × g), and high-molecular-weight DNA was isolated from the cell pellet using the Wizard HMW DNA Extraction Kit (Promega, US) according to the manufacturer’s protocol. DNA quality and length were assessed on Tapestation (Agilent Technologies, US). Nanopore (Oxford Nanopore Technologies, UK) sequencing was performed on unsheared gDNA with no size selection and prepared using Nanopore Q20+ chemistry kit v14 and sequenced on a MinION instrument using an R10.4 (FLO-MIN112) flow cell. The standard ligation protocol for the selected kit was followed with the selection of the long fragment buffer (LFB). Bases were called using Guppy v6.5.7 with the super-accurate (SUP) algorithm with default quality read filtering, generating 300,309 reads and an *N*_50_ read length of 8,763 bp. De novo assembly was performed with Flye v2.9.2 (https://github.com/fenderglass/Flye) (10) followed by contig correction and consensus generation with Medaka v1.8.0 (https://github.com/nanoporetech/medaka). To generate a circular chromosome, overlaps were identified and removed before the assembly was rotated to the gene predicted by prodigal v2.6.3 (11) nearest the middle of the contig with Circlator v1.5.5 (https://github.com/sanger-pathogens/circlator) (12). A closed, circular chromosomal assembly with a genome coverage of 278.0x was produced and annotated using the NCBI Prokaryotic Genome Annotation Pipeline (PGAP) (13). The new assembly is 99.99% (ANI) (14) identical in primary sequence and highly syntenic (15) with the original (Fig. 1). For example, the new assembly and annotation identify 9 ribosomal RNAs, similar to *T. turnerae* strain T7901 (Genbank: GCA_000023025.1, ASM2302v1). However, the new assembly reduces the genome size by 38,994 bp to 5,348,823 bp, contains 56 fewer predicted CDS, and resolves several assembly errors. For all software, default parameters were used except where otherwise noted.

**Figure 1.**
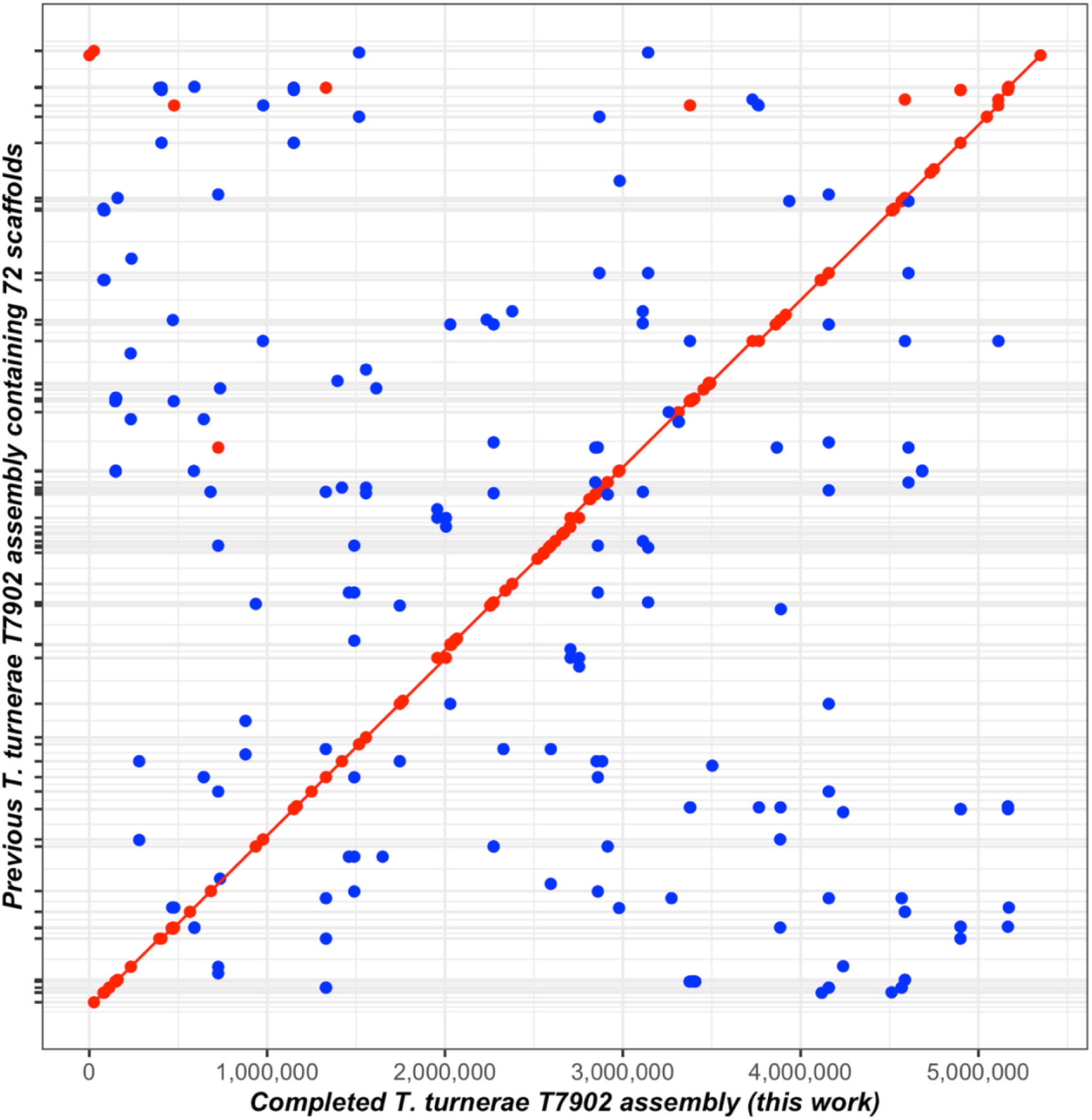
Synteny plot comparing the previously published genome of T. turnerae T7902 (GCA_000379165.1) and the new genome sequence and assembly presented here (GCA_037935975.1). A MUMmer3 plot was generated with NUCmer v3.1 (15) using default settings to assess synteny and completion. Minimum exact matches of 20bp are represented as a dot with lines representing exact match lengths >20bp. Forward matches are displayed in red, while reverse matches are shown in blue.

## Data availability

The complete genome sequence of T7902 has been deposited in GenBank under the accession number CP149817. The Oxford Nanopore sequencing reads are available from the NCBI Sequence Read Archive (SRA) under the accession number SRR28421272.

## Acknowledgments

Research reported in this publication was supported by the following awards to DLD: National Oceanic and Atmospheric Administration (NA19OAR0110303), Gordon and Betty Moore Foundation (GBMF 9339), National Institutes of Health (1R01AI162943-01A1, subaward: 10062083-NE), and Johns Hopkins University Applied Physics Laboratory internal research and development funds. The National Science Foundation (DBI 1722553) also funded some equipment used in this research. The funders had no role in study design, data collection and interpretation, or the decision to submit the work for publication.

